# Serial protein crystallography in an electron microscope

**DOI:** 10.1101/682575

**Authors:** Robert Bücker, Pascal Hogan-Lamarre, Pedram Mehrabi, Eike C. Schulz, Lindsey A. Bultema, Yaroslav Gevorkov, Wolfgang Brehm, Oleksandr Yefanov, Dominik Oberthür, Günther H. Kassier, R. J. Dwayne Miller

## Abstract

Serial X-ray crystallography at free-electron lasers allows to solve biomolecular structures from sub-micron-sized crystals. However, beam time at these facilities is scarce, and involved sample delivery techniques are required. On the other hand, rotation electron diffraction (MicroED) has shown great potential as an alternative means for protein nano-crystallography. Here, we present a method for serial electron diffraction of protein nanocrystals combining the benefits of both approaches. In a scanning transmission electron microscope, crystals randomly dispersed on a sample grid are automatically mapped, and a diffraction pattern at fixed orientation is recorded from each at a high acquisition rate. Dose fractionation ensures minimal radiation damage effects. We demonstrate the method by solving the structure of granulovirus occlusion bodies and lysozyme to resolutions of 1.55 Å and 1.80 Å, respectively. Our method promises to provide rapid structure determination for many classes of materials with minimal sample consumption, using readily available instrumentation.

## Introduction

An understanding of macromolecular structure is crucial for insight into the function of complex biological systems. Despite recent advances in single-particle cryo-electron microscopy (cryo-EM), the vast majority of high-resolution structures are determined by crystallographic methods (http://www.rcsb.org/stats/summary). This includes the majority of membrane proteins, which are often too small for computational alignment as required by single-particle analysis ^1,2^. An important limitation of biomolecular crystallography lies in the difficulty to obtain large, well ordered crystals, which is particularly prevalent for membrane proteins and macromolecular complexes. Sub-micron crystals can be obtained more readily and are a common natural phenomenon, but often escape structure determination as the small diffracting volume and low tolerated dose of typically tens of MGy ^3,4^ prohibit the measurement of sufficient signal. However, during the past few years crystallographic techniques have emerged that are able to exploit nanocrystals for diffraction experiments. Notably, X-ray free-electron lasers (XFELs) have driven the development of *serial* crystallography ^5–10^, a technique that is also increasingly applied at synchrotron sources ^11–17^. Here, acquiring snapshots in a single orientation from each crystal instead of a rotation series avoids dose accumulation, permitting higher fluences which concomitantly decreases the required diffracting volume. Sufficient signal-to-noise and completeness is achieved through merging of many thousands of such snapshots. Ideally, radiation damage effects are entirely evaded either by a “diffract-before-destroy” mode using femtosecond XFEL pulses ^5^ or by imposing doses too low to cause significant structural damage of each crystal, which has also been implemented at synchrotron micro-focus beam lines ^11,12,18^. However, the scarcity and costliness of XFEL beamtime limits the use of protein nanocrystals for routine structure determination. The development of serial crystallography using smaller scale, ideally laboratory-based instrumentation is therefore highly desirable.

Electron microscopes are a comparatively ubiquitous and cost-effective alternative for measuring diffraction from nanocrystals. While the low penetration depth of electrons renders them unsuitable for large three-dimensional crystals, their physical scattering properties are specifically advantageous for sub-micron crystals of radiation-sensitive materials. Compared to X-rays, the obtainable diffraction signal for a given crystal volume and tolerable radiation dose is up to three orders of magnitude larger due to the higher ratio of elastic to inelastic electron scattering events and a much smaller energy deposition per inelastic event ^1,19^. While seminal experiments on 2D crystals ^20^ were restricted to a small class of suitable samples, various successful implementations of 3D rotation electron diffraction (3D ED) solving structures of beam-sensitive small molecules ^21–23^ sparked interest in applying 3D crystallography also to biomolecules, a technique also referred to as MicroED ^24–26^. Several research groups have now succeeded in solving protein structures by merging electron diffraction data from as little as one up to a few sub-micron sized vitrified protein crystals using rotation diffraction ^27–30^, and very recently the first unknown protein structure could be determined ^31^. Automated procedures are becoming increasingly available to reduce the manual effort of identifying suitable crystals, acquiring rotation series while keeping the crystal under the beam, and sequentially addressing many crystals to be merged ^32–36^. However, despite the high dose efficiency of electrons, damage accumulation throughout the rotation series remains a limiting factor, and acquisition as well as sample screening require careful operation at extremely low dose rates ^37^. Recently, a serial electron diffraction (SerialED) scheme has been introduced for small-molecule crystals where, similar to the aforementioned X-ray experiments, still-diffraction snapshots were obtained and used for structure determination ^38^.

Here we apply serial electron diffraction to protein nanocrystals, using a dose-efficient automated data collection scheme that enabled us to solve the highest-resolution protein structure by electron diffraction to date. This method provides a viable alternative to serial femtosecond crystallography for the determination of high-resolution protein structures from sub-micron sized crystals using laboratory-based instrumentation.

## Results

### STEM-based serial electron diffraction

We perform protein crystallography by serial electron diffraction using a parallel nano-beam in a scanning transmission electron microscope (S/TEM). Analogous to the approach of serial X-ray crystallography, we mitigate the problem of damage accumulation by exposing each crystal only once with a high degree of automation and ease of use. A recently developed indexing algorithm ^39^ allows crystal orientation to be determined followed by merging into a full crystallographic data set. Our serial electron diffraction (SerialED) approach operates on crystals randomly dispersed on a TEM grid and consists of two automated steps. First, the sample is moved to a previously unexposed grid region and an arbitrary, fixed goniometer tilt angle is chosen. An overview image of a TEM grid region is recorded in scanning (STEM) mode at a negligible radiation dose (≈ 5% of that later used for diffraction acquisition), and the positions of the crystals are automatically mapped ^32,40^ (Figure 1a). Second, still electron diffraction patterns are recorded from each (randomly oriented) crystal, synchronizing the microscope’s beam deflectors with a high frame rate camera (Figure 1b). No sample rotation is performed. Thereby, a hit fraction approaching 100% with a peak data collection rate of up to thousands of diffraction patterns per second can be achieved. While the former is defined by the accuracy of the mapping algorithm used to identify crystals in the STEM overview image, the latter is limited only by source brightness and camera frame rate. After completion of the diffraction acquisition, the sample is moved to a fresh region, and the sequence is repeated until sufficiently many diffraction patterns have been collected. Importantly, no special sample delivery devices are required, since the full workflow is conducted in a conventional S/TEM or dedicated STEM instrument. The nano-beam diameter can be matched to the typical crystal size of the sample under study by choosing an appropriate condenser aperture and microscope probe mode, minimizing background scattering and diffraction from multiple lattices.

**Figure 1:**
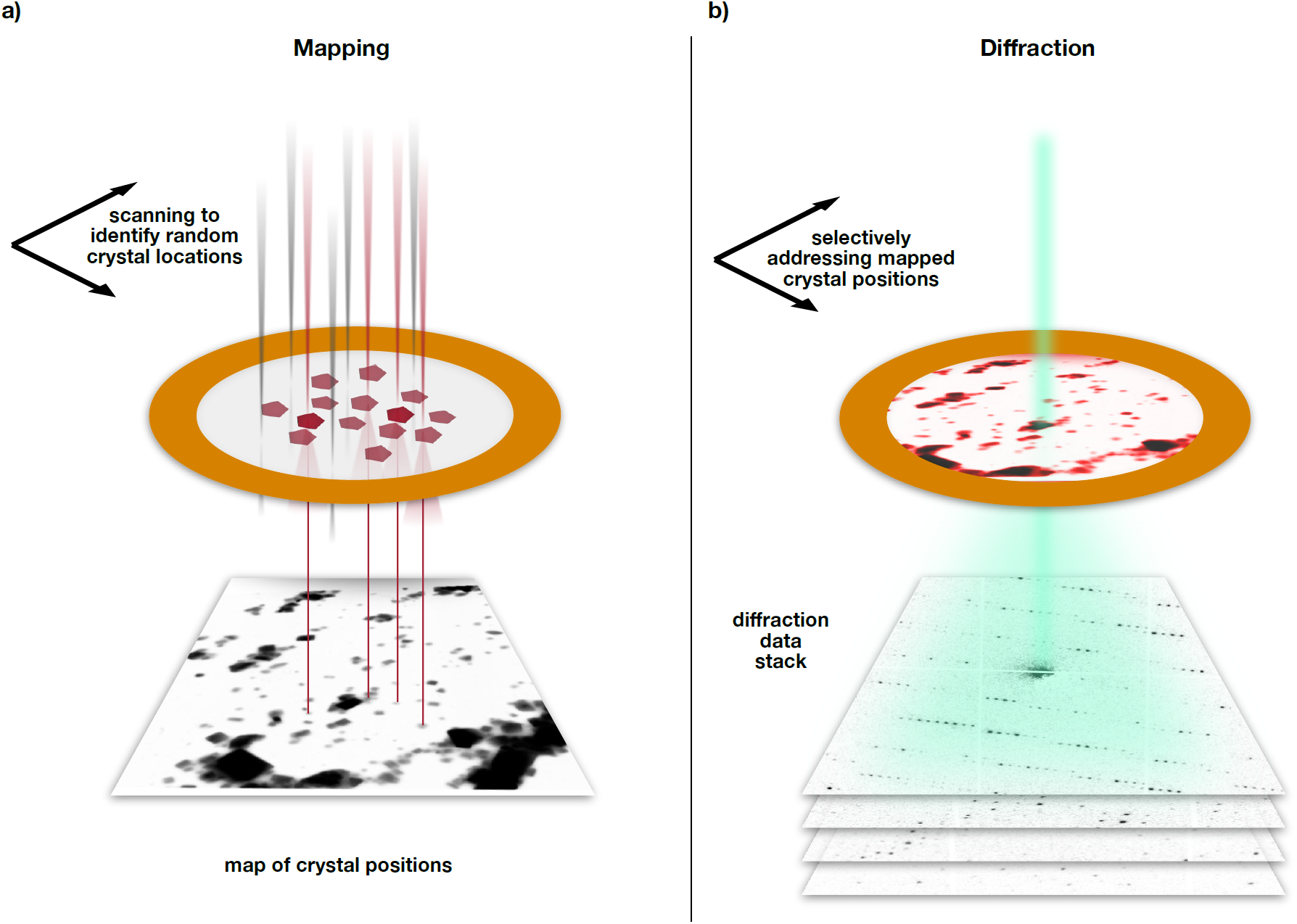
Serial nano-beam electron diffraction scheme. (a) The sample is first mapped in low-dose STEM mode over a large region (typically ≈ 20 µm edge length), yielding a real-space image. Crystals show up as clear features and can be identified automatically. (b) The beam of ≈ 100 nm diameter is sequentially steered to each found crystal position, and diffraction patterns are acquired at a rate of up to 1 kHz.

### Granulovirus occlusion bodies

As a test system, we have chosen natively grown, vitrified granulovirus particles with crystalline occlusion bodies (granulin). The particle size of approximately 100 × 100 × 300 nm^3^ and morphological homogeneity makes this system an ideal target for serial nanocrystallography. Furthermore, granulovirus has previously been studied at LCLS ^7^, and is therefore well suited for purposes of comparing the SerialED approach to XFEL data. We acquired approximately 32000 diffraction patterns from a total sample area of 0.036 mm^2^ on a vitrified TEM grid within a 4 h net measurement duration, that is, including auxiliary steps such as manual search for suitable grid regions containing a large number of viruses embedded in a sufficiently thin ice layer, acquisition of mapping images and automatic crystal identification. Within each grid region, we achieved an average hit fraction of 69% at an acquisition rate of ≈ 50 Hz (see section on dose fractionation below and Supplementary Figure 2). Each crystal was measured in a single orientation, with the goniometer tilt occasionally changed between acquisition runs of different regions (up to 40°) to mitigate effects of preferred sample orientation. Of these hits, 81% could subsequently be indexed and used for merging into a fully complete data set (Supplementary Figure 4). We obtained a complete data set and Coulomb potential maps of excellent quality at 1.55 Å resolution (*R*_free_/*R*_work_ = 0.19/0.17), according to the *CC*\* > 0.5 cut-off criterion ^41^ (Figure 2, Figure 3, Table 1), improving on published XFEL data ^7^ at 2.00 Å resolution.

**Table 1:** Data collection and refinement statistics. Highest-shell values are shown in parentheses.

| Parameter | Granulovirus | Lysozyme |
| --- | --- | --- |
| Electron energy / wavelength | 200 kV / 0.025 Å | 200 kV / 0.025 Å |
| Dose (fluence) per crystal | 4.7 e<sup>-</sup>/Å<sup>2</sup> | 2.6 e<sup>-</sup>/Å<sup>2</sup> |
| Space group | I 2 3 | P 4<sub>3</sub> 2<sub>1</sub> 2 |
| Unit cell |  |  |
| *a*, *b*, *c* | 103.2, 103.2, 103.2 Å | 79.1, 79.1, 38 Å |
| *α*, *β*, *γ* | 90, 90, 90 ° | 90, 90, 90 ° |
| No. of hits / indexed lattices | 20,608 / 17,974 | 1,339 / 1,051 |
| Resolution range | 72.97-1.55 (1.61-1.55) Å | 55.93-1.80 (1.86-1.80) Å |
| Unique reflections | 25,971 (2,577) | 15,284 (927) |
| Multiplicity | 495 (70) | 26 (7) |
| Completeness (%) | 100 (100) | 77.7 (50.4) |
| *R*<sub>split</sub> (%) | 8.7 (168.6) | 16.7 (28.3) |
| Mean *I*/σ(*I*) | 9.22 (0.71) | 6.72 (4.64) |
| CC<sub>1/2</sub> | 1.00 (0.19) | 0.94 (0.22) |
| CC* | 1.00 (0.56) | 0.98 (0.60) |
| Wilson B-factor (Å<sup>2</sup>) | 12.3 | 9.59 |
| Reflections used in refinement | 26,635 (2,630) | 9,067 (569) |
| Reflections used for R-free | 1,297 (149) | 436 (25) |
| *R*<sub>work</sub> (%) | 17.1 | 27.1 (37.2) |
| *R*<sub>free</sub> (%) | 19.7 | 31.6 (38.6) |
| Number of non-hydrogen atoms | 2128 | 1069 |
| macromolecules | 2038 | 1009 |
| solvent | 90 | 60 |
| RMS bonds / angles | 0.012 Å / 1.13 Å | 0.003 Å / 0.47° |
| Ramachandran favored (%) | 97.1 | 96.1 |
| Ramachandran allowed (%) | 2.90 | 3.94 |
| Ramachandran outliers (%) | 0.41 | 0.00 |
| Rotamer outliers (%) | 0.45 | 0.94 |
| Clashscore | 1.74 | 7.58 |
| Average B-factor | 19.3 | 9.5 |
| macromolecules | 19.2 | 9.5 |
| solvent | 20.4 | 9.9 |
| PDB-ID | 6S2O | 6S2N |

**Figure 2:**
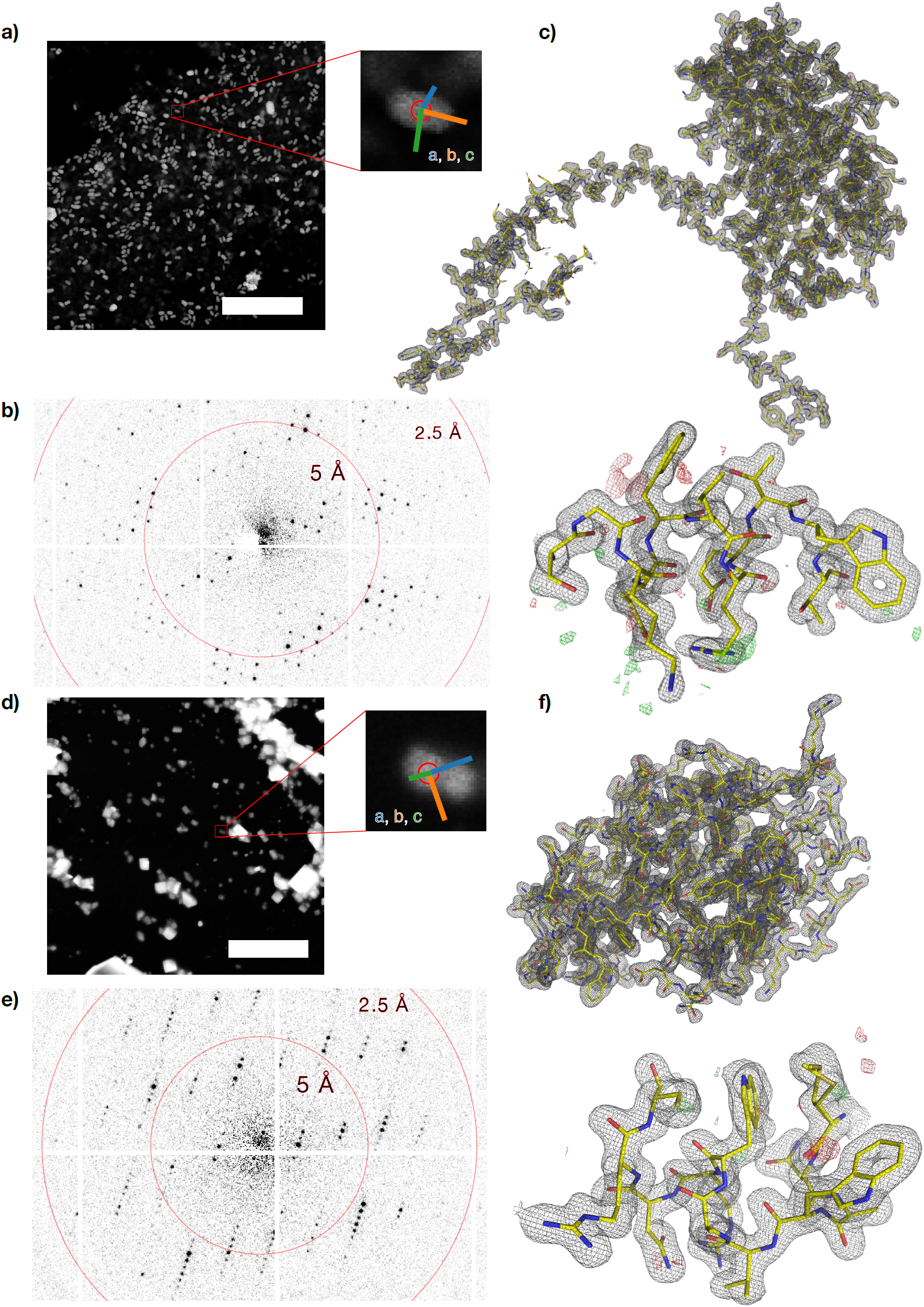
SerialED results for granulovirus occlusion bodies and lysozyme. (a) STEM mapping image of a grid section containing granuloviruses, visible as bright features (scale bar is 5 μm). A zoomed view of a representative virus is shown, where the red circle corresponds to the diffraction nano-beam diameter of ≈110 nm. Coloured lines indicate the lattice vectors found after indexing of the diffraction pattern. (b) Diffraction pattern acquired from the features shown in (a). (c) Obtained structures of granulin; 2F_o_-F_c_ map of the entire structure, and zoom into a randomly chosen region, with F_o_-F_c_ map overlaid. Both maps are at 1.55 Å resolution and contoured at ±1σ. (d-f) Analogous for lysozyme nanocrystals; maps are at 1.8 Å resolution.

**Figure 3:**
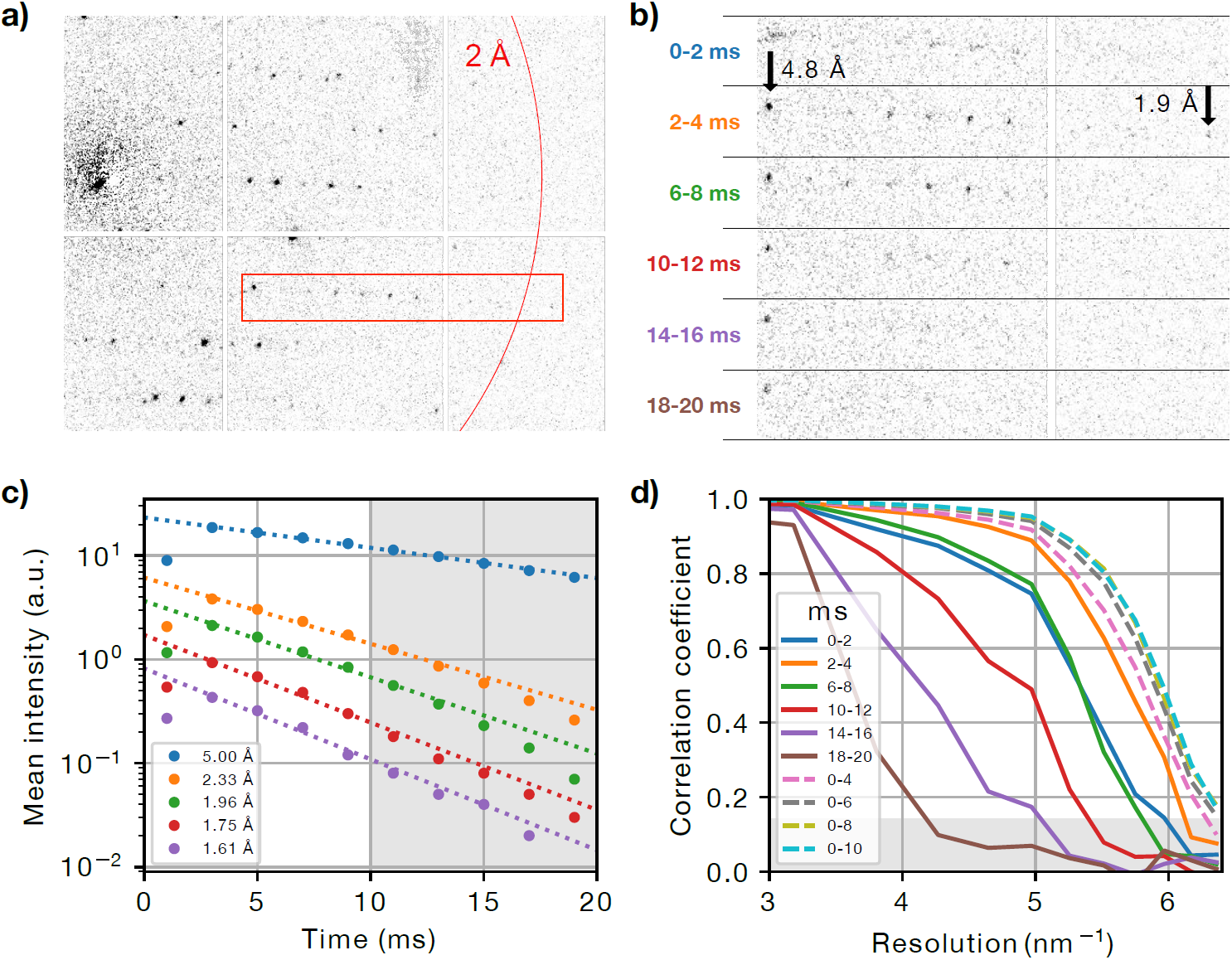
Radiation damage during dose-fractionated acquisition. (a) Typical diffraction pattern from a granulovirus occlusion body. The red box indicates the enlarged region in b). (b) Enlarged diffraction pattern section for several single frames from the dose-fractionated movie stack, each of 2 ms duration. The integration time of each frame relative to the beam first hitting the crystal is specified. Note the fading of the diffraction spots especially at high resolutions. The first shot is affected by residual beam motion and hence has a shorter effective integration time and shows blurring artefacts. (c) Mean intensity of Bragg reflections for different resolution shells as a function of delay time, and exponential fit lines, where the first time point has been excluded from the fit. The shaded area corresponds to delay times beyond 10 ms, which have been excluded from our data analysis. (d) Resolution-dependent correlation coefficients *CC*_1/2_ shown from 3.33 Å to Å resolution. Solid lines correspond to single movie frames as in (b). Dashed lines correspond to images that were cumulatively summed over several frames. The shaded area corresponds to values *CC*_1/2_ < 0.143, where data falls below the resolution cut-off at *CC*\* = 0.5.

### Lysozyme

Furthermore, we applied the SerialED method to the common test sample hen egg-white lysozyme (HEWL). HEWL crystals of typically 100-500 nm diameter were deposited on a standard TEM-grid and vitrified (see Methods). Two independently prepared samples were measured in separate acquisition runs over a total measurement duration of 3 h and a sample area of 0.010 mm^2^. Diffraction patterns from 1325 nanocrystals were collected, achieving a hit fraction of 62% at an acquisition rate of 50 Hz. 83% of the obtained patterns could be successfully indexed and used for merging (78% completeness, see Supplementary Figure 4), resulting in a Coulomb-potential map of high quality to 1.8 Å resolution (Figure 2; Table 1). The determined HEWL structure compares well to previously determined structures by X-ray and micro-electron diffraction techniques (see Methods).

### Radiation damage and dose fractionation

The high frame rate and zero background of the detector applied in our experiment allows recording a burst-series comprising several frames instead of a single snapshot for each crystal, yielding a dose-fractionated *diffraction-during-destruction* data stack. Both data sets shown were acquired with the camera running continuously at 500 frames per second; each crystal was exposed for 10 movie frames with the beam resting at the crystal position as determined in the mapping step, resulting in a net acquisition rate of ≈ 50 Hz (Supplementary Figure 2). Therefore, the per-crystal exposure time of 20 ms was fractionated into a stack of diffraction patterns of 2 ms exposure time each, which exhibit a pronounced fading of high-resolution peaks (Figure 3). A final set of diffraction images was generated by cumulatively summing movie frames in the acquired data. Thus, the effective integration time and dose per crystal can be chosen *after* data acquisition has concluded, trading off between low radiation damage (short integration) and high signal-to-noise ratio (long integration). Hence, *a priori* knowledge of the sample’s radiation sensitivity (critical dose) is not required, and data can be obtained before the onset of observable radiation damage. For our data sets, we find an instantly detectable loss of high-resolution Bragg peaks, in accordance with previous studies ^37^ (Figure 3c). In Figure 4c, mean reflection intensities from the granulin data set are shown for several resolution shells. Exponential fits to the data show a fair agreement and lead to estimated 1/e decay times of 14.9(4) ms at 5.00 Å, 6.8(3) ms at 2.33 Å, 5.9(3) ms at 1.96 Å, 5.2(2) ms at 1.75 Å, and 5.0(3) ms at 1.61 Å, the latter corresponding to an approximate dose of ≈ 2.6 e^-^/Å^2^. The optimal integrated dose was found by observing the half-set correlation coefficient *CC*_1/2_ 41 calculated for merged data sets that were derived from diffraction patterns summed over different numbers of movie frames (Figure 3d). For granulin, optimal data quality was reached for summation of the first five movie frames, corresponding to an exposure time of 10 ms, and an integrated dose of ≈ 4.7 e^-^/Å^2^; for lysozyme, we found an optimal dose of ≈ 2.6 e^-^/Å^2^. More detailed measurements of site-specific and global radiation-damage effects, as well as optimization of data acquisition and analysis strategies to further improve dose efficiency and resolution will be the subject of future work.

**Figure 4:**
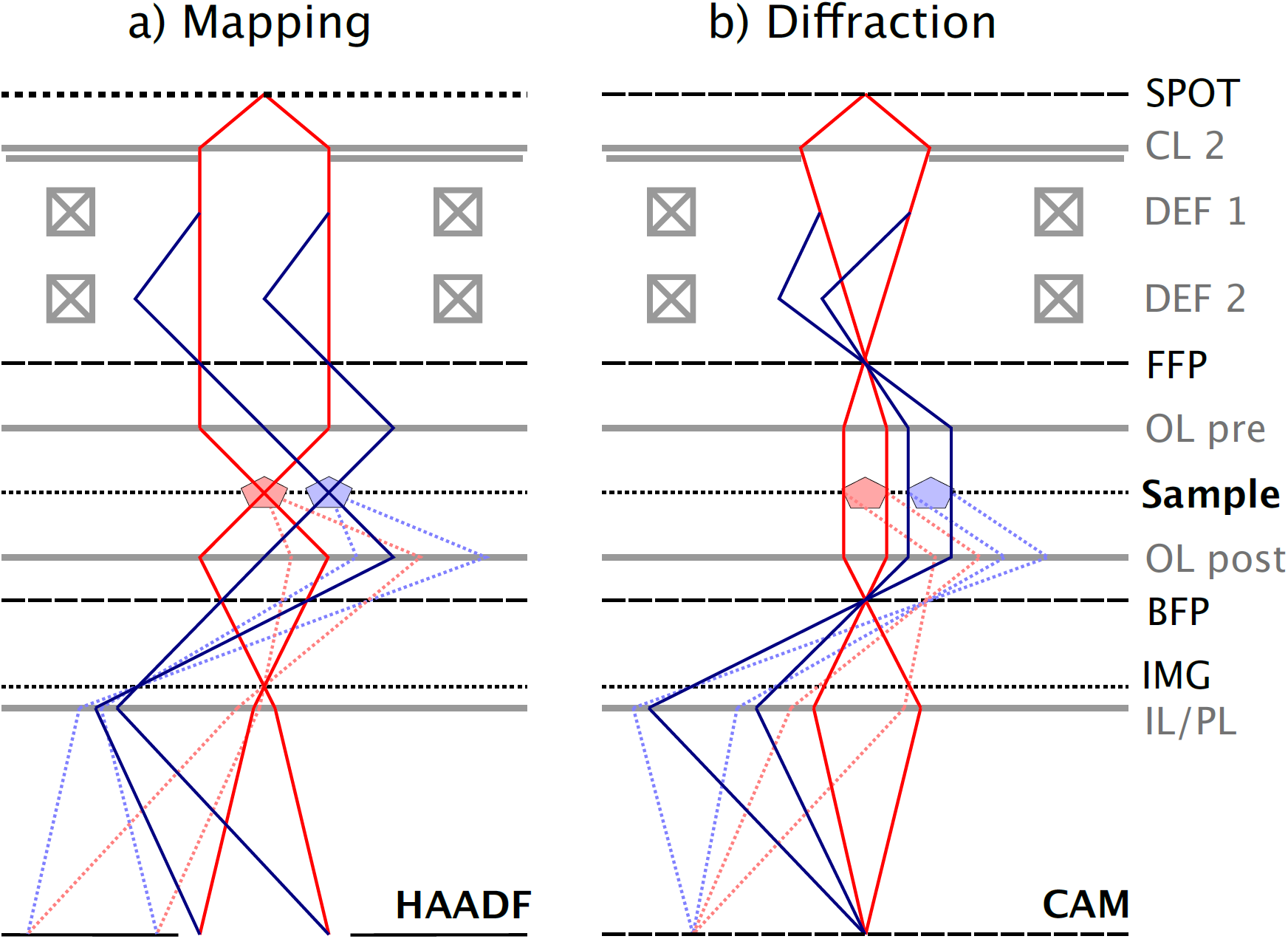
Ray path diagrams for mapping (focused) and diffraction (collimated) condenser configuration. Red and blue lines correspond to on-axis and one exemplary off-axis positions of the beam. Dotted lines correspond to a Bragg reflection. Optical planes and electron-optical elements are shown in black and grey, respectively. (a) In the mapping configuration, the beam is collimated by the lower condenser lens (CL 2) and focused on the sample using the objective lens pre-field (OL pre). Scattered beams from the illuminated sample position are imaged on the high-angle annular dark field (HAADF) detector using the objective lens post-field (OL post) and the intermediate and projection lenses (IL/PL). In the diffraction configuration, on the other hand, the condenser focuses the beam on the front-focal plane (FFP) of the objective. Diffraction orders now appear as discrete spots on the diffraction detector (CAM). Note that switching between these configurations involves changing of the CL 2 excitation only, as the detectors always remain in a plane conjugate with the back and front focal planes of the objective lens (diffraction mode). (SPOT – first condenser lens (spot) crossover; DEF1/2 – upper and lower beam deflector pair; IMG – intermediate image plane).

## Discussion

Our results show that serial electron diffraction (SerialED) allows the determination of protein structures at high resolution from extremely small protein crystals in a rapid, efficient, and automated manner. No sample rotation during measurement of each crystal is required, simplifying the measurement and allowing the use of higher doses for each diffraction pattern. Also, no manual screening and selection of individual suitable crystals under low-dose conditions are necessary. In contrast to wide-field TEM-based crystal mapping ^36,38^, our STEM-based scheme neither requires frequent mode-switching of the microscope (which always remains in diffraction mode) nor accurate beam-position calibrations, as crystal mapping and nano-beam positioning are achieved with the same set of deflectors. Furthermore, the acquisition speed is not limited by relatively slow software-based scripting of the microscope, which is entirely bypassed (see Methods), and a small condenser aperture can be used at all times, achieving a fully parallel (Köhler) nano-beam illumination (Figure 4), thus enabling distortion-free diffraction. Minimal-damage acquisition is ensured using a dose-fractionated diffraction-during-destruction scheme and a-posteriori critical dose determination. We have demonstrated a net acquisition rate of 35 Hz when factoring in the hit fraction (Supplementary Figure 2), which is comparable to current liquid-jet XFEL ^8,9^ and synchrotron fixed-target ^11,12,18^ experiments. Note that a further increase of more than an order of magnitude can be achieved if dose-fractionation is omitted and acquisition speed becomes detector frame rate limited. A complication of SerialED data analysis is the difficulty of determining space group and lattice parameters from single high-energy electron diffraction patterns due to the flatness of the Ewald sphere (λ = 0.025 Å); successful indexing as demonstrated here requires prior knowledge of the crystal parameters. However, those can for instance be determined from an auxiliary low-resolution rotation diffraction data set obtained from few crystals on the same sample. Alternatively, an approach of clustering spot-distance data from *all* acquired patterns and deriving lattice parameters from comparison to forward-modelling has yielded promising results ^42^. Preferred crystal orientation limiting data set completeness is often encountered in electron diffraction ^43^, even when merging a moderately large number of rotation diffraction data sets ^29^. In SerialED, with varying rotation angles as described above, the much higher number of merged crystals which may occasionally assume unusual orientations can lead to a mitigation of this issue (see Supplementary Discussion). A further improvement would be achieved using specialized TEM grids ^44^, or microfabricated chips ^45^. The SerialED approach could also be applied to heterogeneous systems with extended amorphous regions, such as cells containing *in-vivo* grown nanocrystals ^3^, or to map and exploit local lattice structures ^46^. Similarly, mixtures of crystals within a single grid or contaminated samples can be studied without significant modifications by assigning each found lattice to one of the contained sample classes using multiple indexing runs or direct classification of diffraction patterns ^36,47^. It is moreover not only limited to proteins but encompasses all nanocrystalline compounds, such as pharmaceuticals ^47,48^ or porous materials ^23,49,50^. Augmenting parallel-beam crystallography with coherent scanning diffraction techniques such as convergent-beam diffraction or low-dose ptychography might be a viable way to obtain Bragg reflection phase information ^51,52^. Finally, integrating the serial acquisition approach with emerging methods of *in-situ* and time-resolved electron microscopy ^53–55^ may open up avenues for room-temperature structures or structural dynamics studies on beam-sensitive systems. All of this makes STEM-based serial electron diffraction a versatile, highly efficient and low-cost alternative to canonical structure determination approaches for proteins and beyond.

## Methods

### Sample preparation

Commercially available *cydia pomonella* granulovirus of formulation Madex Max was obtained from Andermatt Biocontrol. The occlusion bodies (OBs) were purified from the aqueous suspension by iterative washing and centrifugation cycles. The pellet was then re-suspended in ultra-pure water at pH 7 and subjected to filtration steps through a sequence of nylon mesh filters with decreasing mesh diameter (100 µm, 50 µm, 20 μm, 10 μm, 5 μm, all Sysmex, Germany) and finally twice through 0.5 μm stainless steel filters (IDEX, USA). To increase the concentration of OBs, the suspension was subjected to centrifugation at 21,000 g, and 90% of the supernatant removed.

Hen Egg-White Lysozyme (HEWL) was purchased from Sigma-Aldrich as a lyophilized powder. It was dissolved in 20 mM NaAcetate pH 4.7 to a concentration of 80 mg/ml. HEWL crystals were grown via batch crystallization, whereby equal volumes of the protein solution and 80 mg/ml NaCl were added.

Crystals ranging from 5-10 µm rapidly formed within 2-3 hours. The resulting crystal mixture was centrifuged down and 75% of the supernatant was removed creating a dense crystal slurry. Subsequent vortexing with steel beads in a microfuge tube for 30 minutes resulted in a concentrated suspension of crystal fragments in the sub-500 nm size range.

For each of the above suspensions, 2 µl were applied to 400-mesh carbon grids (type S160-4 purchased from Plano GmbH), whereupon blotting and vitrification using a mixture of liquid ethane/propane was performed in a Vitrobot Mark IV (Thermo Fisher Scientific).

### Diffraction data acquisition

All data has been acquired on a Philips Tecnai F20 TWIN S/TEM, equipped with a Gatan 626 cryo-transfer holder, a X-Spectrum Lambda 750k pixel array detector based on a 6×2 Medipix3 array ^25^, and a custom-built arbitrary pattern generator addressing the deflector coil drivers, based on National Instruments hardware (see Supplementary Methods for discussion of hardware requirements). Initially, the grids were screened in low-magnification STEM mode for regions exhibiting a high crystal density without excessive overlap and aggregation. While software such as SerialEM ^56^ could be used in a straightforward manner to automate this screening step, this was not required for our test samples, as sufficiently homogeneous regions were readily found.

After screening, the microscope was set to standard STEM mode at the lowest possible magnification, corresponding to a (18*μm*)^2^ field of view. To achieve a high-current (≈ 0.1 nA), small (≈ 110 nm), and collimated (≪ 0.5 mrad) nano-beam, the following microscope settings were made: field-emission gun parameters at weakest excitation of both gun lens and C1 condenser lens (Spot size), disabled mini-condenser (nanoprobe mode), small (5 µm) condenser (C2) aperture. The microscope remains in diffraction mode at all times, that is, the back-focal plane of the objective lens is conjugate with the detector.

At each of the identified sample regions, the two-step acquisition sequence as shown in Figure 1 was performed:

1. The beam was focused on the sample (Figure 4a), and an overview STEM image of 1024×1024 pixel resolution was taken across the entire field of view (Figure 1a) using the high-angle annular dark field (HAADF) detector. The dwell time was set such that the exposure dose remained small, well below 0.1 e^-^/Å^2^. From this image, crystals were automatically identified using standard feature extraction methods, and a list of scan points, corresponding to discrete values of the microscope’s scan coil currents, was derived (see Supplementary Methods and Supplementary Figure 1).
2. The beam was defocused into a collimated nano-beam (Köhler illumination) of 110 nm diameter^40^, yielding sharp diffraction patterns in the objective back-focal plane and on the detector (Figure 4b). The actual diffraction data acquisition was then performed by sequentially moving the beam to each of the crystal coordinates using the STEM deflectors and recording a diffraction movie (dose-fractionated data stack) at each position (Figure 1b, Figure 3).

Once data acquisition from the mapping region was complete, the beam was blanked, and the sample stage moved to the next previously identified sample region. This sequence was repeated until several thousand diffraction patterns had been recorded. All steps were automated and controlled using Jupyter notebooks based on Python 3.6, and a custom instrument control library written in LabVIEW and Python 3.6, using parts of the *Instamatic* library ^38^.

### Data processing

The recorded diffraction patterns were pre-processed using our *diffractem* package (www.github.com/robertbuecker/diffractem), setting up a pipeline comprising dead-pixel and flat-field detector correction, partial summing of dose-fractionation stacks, as well as centring of each pattern using the position of the transmitted beam and position-matching of simultaneously excited Friedel-mate reflections (Supplementary Figure 3). Diffraction spots were identified using the peakfinder8 algorithm contained in the *CrystFEL* suite ^57,58^; patterns containing more than 25 spots at resolutions below ≈ 2.5 Å were selected for further analysis. The extracted spot positions were used to determine the orientation of each crystal, and to predict the position of the corresponding Bragg reflections using the indexing and refinement algorithm *PinkIndexer* ^39^. Intensities of the Bragg reflections were extracted using two-dimensional peak fitting, and a full reciprocal-space data set was obtained using the *partialator* program from *CrystFEL* ^59^. Data were truncated after the last resolution shell where *CC\** > 0.5 ^41,60^. Refer to the Supplementary Methods for additional details on data pre-processing, indexing, and merging. Phasing of the models was achieved by molecular replacement using *Phaser* ^61^ from the *PHENIX* software suite ^62^ using PDB-ID: 4ET8 and PDB-ID: 5G3X as template models, respectively. Upon obtaining phases, iterative cycles of model building were made using *Coot* ^63^. For correct refinement of the Coulomb potential maps, subsequent rounds of refinement were performed using *phenix.refine*, taking electron scattering factors into account ^64^. Illustrations of the electron density map and model were generated using PyMOL by Schrödinger. Crystallographic statistics are reported in Table 1. To validate the consistency of our structures with known data, we calculate the rms deviations (RMSD) of atom positions with respect to previously published structures. For Lys with respect to PDB-ID 5K7O (measured by MicroED) we find a value of RMSD = 0.487 Å; for Lys w.r.t. 5WR9 (measured by XFEL serial crystallography) RMSD = 0.353 Å. For GV with respect to PDB-ID 5G3X (synchrotron crystallography) we find RMSD = 0.206 Å; for GV w.r.t. 5G0Z (XFEL) RMSD = 0.353 Å.

## Data availability

An example sub-set of raw experimental data is available from the MPG Open Access Data Repository at [https://dx.doi.org/10.17617/3.2j]. The full raw data sets are available from R.B. upon request. CrystFEL *stream* and *hkl* files containing unmerged and merged reflection intensities, respectively, are available as Supplementary Material to this manuscript. The protein structures can be accessed from wwPDB using the codes 6S2O [http://dx.doi.org/10.2210/pdb6S2O/pdb] (Granulin) and 6S2N [http://dx.doi.org/10.2210/pdb6S2N/pdb] (Lysozyme).

## Code availability

*PinkIndexer* and *CrystFEL* are available at http://www.desy.de/~twhite/crystfel/ under the terms of the GNU general public license. *diffractem* is available at www.github.com/robertbuecker/diffractem under the terms of the MIT license. Python-based Jupyter notebooks executing the data analysis pipeline using *diffractem* and *CrystFEL* are available along with the example raw data from the MPG Open Access Data Repository at https://dx.doi.org/10.17617/3.2j All custom software for data acquisition and hardware control is available from R.B. upon request.

## Acknowledgments

We thank Djordje Gitaric for mechanical design work, Fabian Westermeier, David Pennicard, and Heinz Graafsma for adapting the *Lambda* detector for electron imaging, Anton Barty and Henry Chapman for many helpful discussions and critical reading of the manuscript, Thomas A. White for help with modifying *CrystFEL* for electron diffraction, and Michiel de Kock for help with image processing. We are indebted to Kay Grünewald and his research group for lending to us their cryo-transfer holder. We gratefully acknowledge the support provided by the Max Planck Society, the excellence cluster ‘The Hamburg Centre for Ultrafast Imaging—Structure, Dynamics and Control of Matter at the Atomic Scale’ of the Deutsche Forschungsgemeinschaft EXC 1074 project ID 194651731 (R.J.D.M.), and the Joachim Herz Foundation (Biomedical Physics of Infection). P.M. was supported by the Alexander von Humboldt-Stiftung for postdoctoral researchers.

## Author contributions

R.B., G.K. and R.J.D.M. conceived the SerialED scheme. R.B. implemented the experimental set-up and conducted the measurements. P.M. and G.K. prepared the lysozyme nanocrystals, D.O. prepared the granulovirus sample, and G.K. and L.B. prepared vitrified cryo-EM grids from each sample. R.B., P.H. and P.M. performed the data processing. Y.G., O.Y. and W.B. adapted the *PinkIndexer* code to electron diffraction. R.B., E.S. and G.K. wrote the paper, with contributions from all authors.

## Supplementary Methods

### Instrumentation

The serial electron nano-beam diffraction scheme can in principle be performed in any S/TEM or dedicated STEM with a sufficiently fast, hardware-synchronizable camera and software scripting interface. We used a Philips Tecnai F20 S/TEM with a TWIN pole piece, a Schottky field-emission gun, a Fischione Model 3000 HAADF-STEM detector, and a X-Spectrum Lambda 750k camera.

For beam currents in the tens of picoamperes range, the camera frame rate should be at least tens of Hertz so as not to limit the acquisition throughput. Implementation of the dose-fractionated movie mode requires a significantly higher frame rate, ideally hundreds of Hertz or greater. A hybrid pixel detector ^25^ meets these requirements optimally, as long as the count rates do not exceed the saturation threshold. The camera used in our work was a 6×2-panel Medipix3-based detector operating in 12-bit continuous-readout mode at up to 2 kHz and a resolution of 1536×512 pixels. Scintillator-coupled detectors based on latest-generation fast CMOS sensors are a viable and commonly available alternative. Back-thinned monolithic pixel detectors as often used in cryo-EM imaging have also been reported to provide sufficient radiation-hardness and dynamic range ^65^.

To direct the sequential beam motion, the X/Y (line/frame) control voltages of the STEM deflector drivers were addressed from an off-the-shelf PC-based data acquisition board (National Instruments PCI-6251). The list of scan points derived as described below is directly written into the output buffer of the board’s digital-analogue converters. While the data acquisition is running, synchronized trigger signals for the camera are provided. The same hardware is also used for acquiring data from the HAADF-STEM detector during the mapping step.

To control the microscope, detector, and scan generator, we use custom software based on Python 3.6 and National Instruments LabVIEW, implementing high-level functions for serial crystallography workflow automation (available from R.B. on request). Instead of a dedicated graphical user interface, we use *Jupyter* notebooks to control an acquisition run and visualize its progress, which can be adjusted to the sample under study and annotated, providing a self-documenting protocol for each data acquisition.

### Instrument preparation

In the following, we lay out in more detail the steps required to acquire a serial crystallography data set as performed in our work. Before executing the procedure outlined in the methods section, the following preparation steps have to be taken:

- Common parameters of the microscope should be properly aligned and characterized. Of specific importance are gun tilt and shift, STEM pivot points, stigmators, rotation center (beam tilt), and centring of the condenser (C2) aperture. The camera length and any diffraction distortion should be carefully calibrated using a standard polycrystalline target such as thallous chloride (TlCl). While helpful for interpreting the real-space (STEM) maps, a precise calibration of the deflectors (STEM magnification and distortion, static beam shift) is explicitly not required.
- By changing the settings of the electron gun (spot size, gun lens, and extraction voltage on a FEG instrument; spot size, filament current and Wehnelt bias on a thermionic instrument), the beam current is optimized to match the beam diameter *d*, camera frame rate *f*, desired total dose (fluence) *D*, and number of dose-fractionation movie frames *K* as *I* = *eDfπ*(*d*/2)^2^/*K*, where *e* denotes the elementary charge.
- The setting of the condenser lens corresponding to a collimated beam is determined by focusing diffraction spots with the projection system in diffraction mode and the diffraction lens focused on the back focal plane of the objective lens (Diffraction setting, Figure 4b). Diffraction data is taken using this setting, as described in the main text.
- The sample is brought to the eucentric height by minimizing image motion when wobbling the stage tilt. The condenser setting (defocus in STEM mode) required to achieve a focused STEM image is noted or stored in the automation software (Mapping setting, Figure 4a). Now, the position of the condenser aperture (C2) can be precisely aligned by switching the microscope repeatedly between focused (mapping) and collimated (diffraction) condenser lens settings and observing the position of the beam in the sample plane ^40^. In our microscope we notice that the residual beam shift between both settings can be minimized by renormalizing the illumination system after each change between settings. Even without renormalization, a satisfactory repeatability can however be reached, as long as the condenser lens is not intermittently set to other values.
- Finally, the offset of the STEM mapping image along the fast-scanning axis *x* due to finite scan speed Δ*x*_scan_ needs to be determined. This can be achieved by recording STEM data along a few discrete *y* coordinates only but spanning the full range of the *x*-axis. This is repeated twice, once for the scan parameters as used for the mapping image (dwell time, pixel size, resolution, magnification, fly-back time, detector time constant, etc.), and once at a low scan speed (dwell time of typically 1 ms, no fly-back time), with the beam assuming a quasi-static position within each pixel dwell time. Δ*x*_scan_ can be determined via a straightforward cross-correlation registration between the obtained intensity data. The offset calibration is fully automated and requires less than one minute. We find that the obtained value remains stable over measurement sessions spanning several days.

**Supplementary Figure 1:**
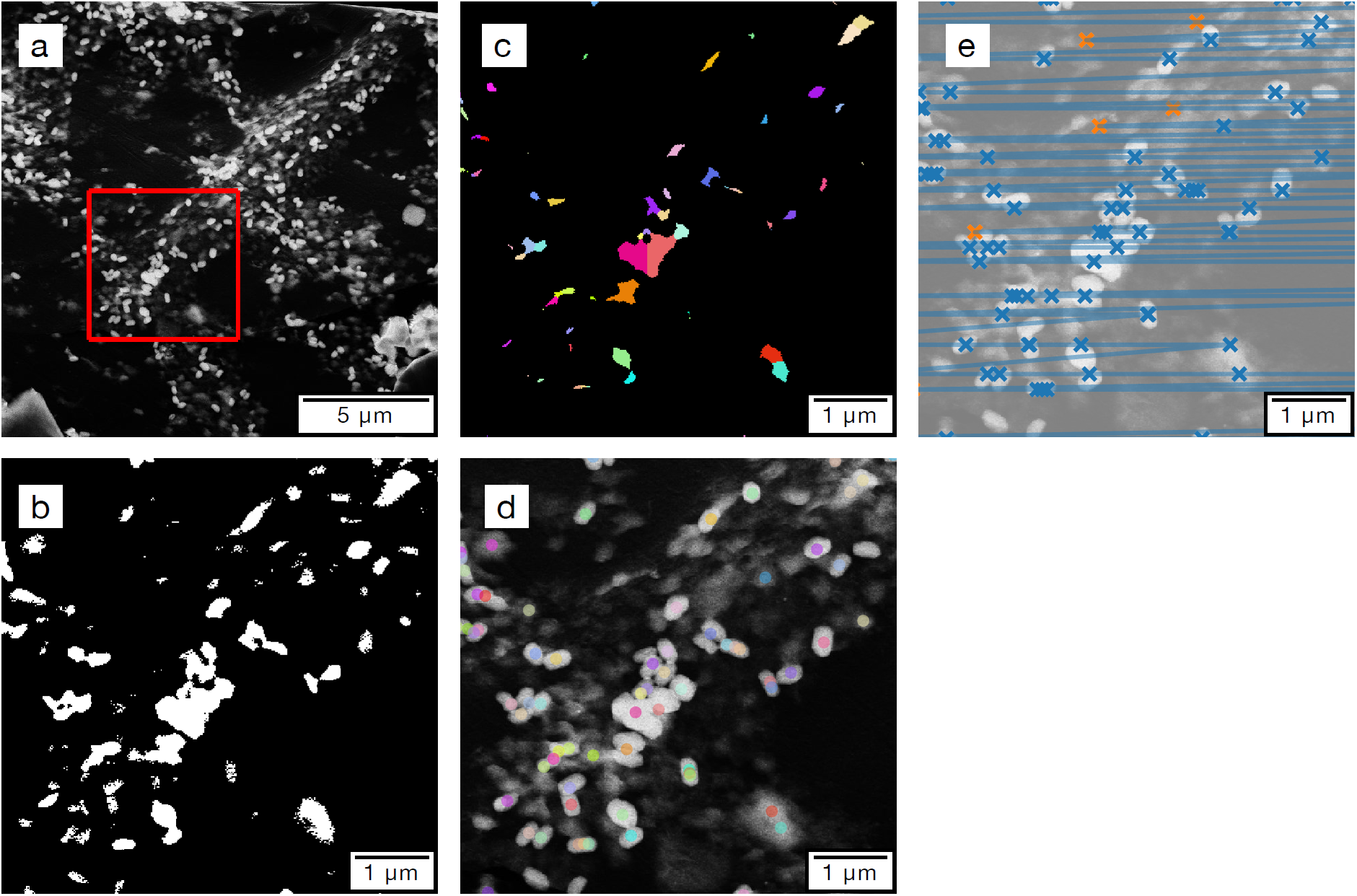
Particle picking and scan path generation. (a) Typical mapping image, with granuloviruses and residual solvent contamination. The red square indicates the zoomed region of panels (b-e). (b) Zoomed view after binarization using an automatically determined threshold. (c) Labelled regions after segmentation. (d) Mapping image with final beam coordinates for diffraction. (e) Scan path with acquisition points (blue) and auxiliary points (orange).

### Crystal finding and acquisition programming

Once the mapping image has been acquired using STEM, the coordinates of the nano-beam for diffraction recording, corresponding to crystal features, are determined using the following automated image analysis procedure, which can be adapted in its parameters to the sample of interest:

- The image is binarized using a fixed or automatically determined grey-value threshold, the latter derived using Otsu’s or Li’s method ^66,67^.
- A morphological closing operation with a structuring element of a typical minimum size of a single crystal is performed to exclude noise and features that are too small.
- Crystals are often found within aggregates or thicker sample regions, which are registered as a single bright segment. A second round of thresholding and binarization can now be performed on each individual bright segment to locate individual crystals.
- In order to further discern crystals in connected regions, a watershed segmentation starting from either local intensity or distance-transform maxima is performed. The resulting segments are assumed to belong to a single crystal and assigned a unique ID.
- For each segment, a diffraction beam position is selected by determining its centre of mass. Alternatively, multiple beam positions spaced by a distance of approximately the nano-beam diameter can be distributed over the segment, using a k-means clustering approach similar to ^38^. While this has not been performed in the present work, it may prove beneficial for cases where clear boundaries of adjacent crystals are not readily discernible, or to study and exploit local lattice structures ^46,68^.

**Supplementary Figure 2:**
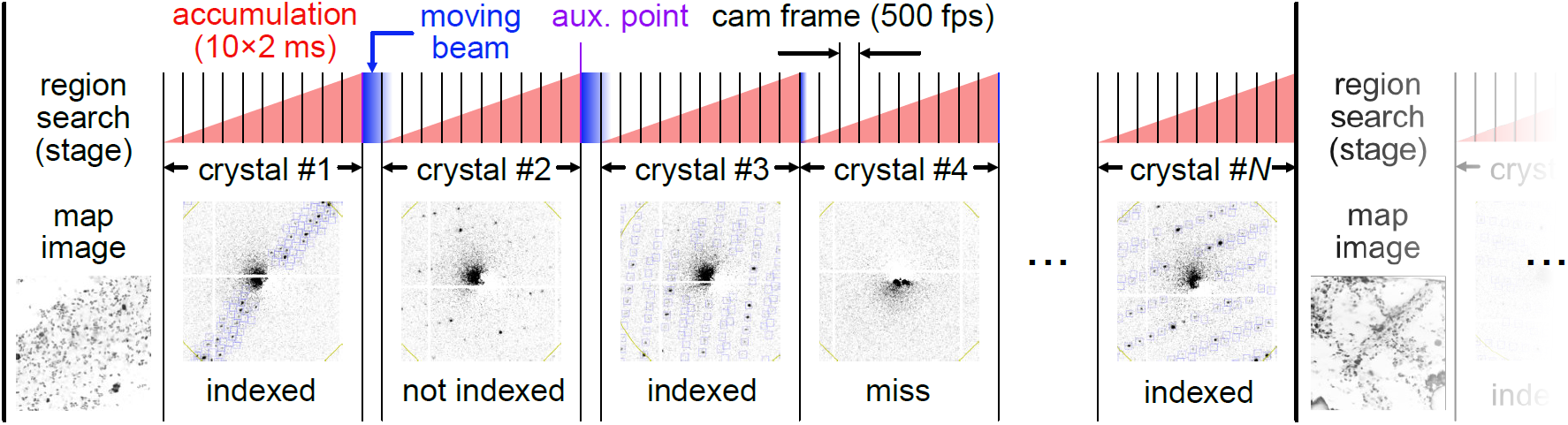
Acquisition timing structure. Left to right: first, a suitable region for data collection on the sample grid is identified, and crystal positions are mapped. Then, the acquisition sequence is started, during which the camera is running at a fixed frame rate (here: 500 frames per second), and a dose fractionation stack from each crystal is recorded for a given number of frames (here: 10), which can contain usable diffraction data (“indexed”), no data (“miss”) or unindexable diffraction data (“not indexed”). Next, if required, an auxiliary scan point is inserted as described in the text, and the beam is quickly moved to the next crystal. In our example, data from slightly less than 50 beam positions per second are recorded, corresponding to indexed patterns at a rate of ≈ 35 Hz. Afterwards, the sequence is repeated at a fresh sample region.

At this point, the coordinates of the desired beam positions for recording diffraction patterns are known. Note that these coordinates do not have to be calibrated to real space, but merely represent control voltages of the deflection coil drivers applied during the mapping image acquisition. Next, a list of scan points, by which we mean the nominal values of the scan generator outputs for the diffraction acquisition step, are derived. Due to effects such as beam hysteresis, these have to be corrected and modified with respect to the previously determined crystal coordinates. The derivation is conducted as follows:

- The crystal coordinates of all crystals along the *y* axis (vertical, slow-scanning) are clustered into a discrete set (scan rows at coordinates *y*’) using a one-dimensional k-means algorithm. The number of scan rows is lower than the total number of crystals but chosen such that the maximum deviation between desired (*y*) and discretized (*y*’) coordinates remains below a given threshold, typically chosen as half the scanning beam radius. Coordinates along the *x* axis (horizontal, fast-scanning) remain unaffected. At this point, the list of scan points is initialized from the (*x, y*′) coordinate tuples.
- Scan points are identified, where the distance to the previous one along either axis is either negative or exceeding an empirically determined threshold. Before each such point, an auxiliary scan point is inserted, at the same *y*’-position as the actual point, and an *x*-position reduced by a certain amount. The dwell time at the auxiliary points is typically shorter than on the actual recording points, and either no diffraction data is recorded for them, or it is discarded in later processing steps. This step ensures that each crystal is approached from the same direction (from the top and/or from the left) from a distance that is not exceedingly large. While the former ensures position reproducibility despite lens hysteresis, the latter helps to avoid artefacts in the diffraction patterns arising from the finite beam scan speed.
- Finally, the offset Δ*x*_scan_ obtained from calibration procedures as described above are applied.

The full algorithm to derive a list of scan points from a STEM mapping image is illustrated in Supplementary Figure 1. The obtained list of scan points is then written into the memory of the scan generator. Dose-fractionation movies are implemented as repeated points with identical coordinates, each one triggering a new camera acquisition. In Supplementary Figure 2, the timing structure of a data collection sequence generated using this prescription is shown. For our example data sets, an effective hit rate of indexable crystals of ≈ 35 Hz is reached. Factoring in the auxiliary steps (region search, map acquisition, sample handling), the net rate reduces to approximately 75 patterns per minute.

### Data pre-processing

We now describe the pre-processing protocol which can be carried out using our *diffractem* software package. Raw detector data for each single acquisition run is initially contained in HDF5 files according to the *NeXus* specification ^69^, which is commonly used in X-ray diffraction. The diffraction data is arranged in a three-dimensional image stack, with a height *Kn*_cryst_ + *n*_aux_, with *n*_cryst_ the number of crystals in the sample region, *K* the number of dose-fractionation movie frames, and *n*_aux_ the number of auxiliary points inserted as described above. Furthermore, the scan position list generated as outlined before, the mapping STEM image with metadata for each found feature, and all accessible settings of the microscope, detector, and scanning unit, are stored within the *NeXus* file.

Starting with these raw input files, the steps outlined in the following are performed to pre-process the data set for use in standard diffraction data reduction software. Using the Python package *dask*, all operations are performed using chunked lazy evaluation in a single calculation step, and efficiently scale on multi-processor systems, with only modest memory requirements; metadata are handled using the *pandas* package.

- The recorded diffraction data are filtered such that all images corresponding to auxiliary scan points are removed. If dose-fractionation movies have been recorded, an effective integration time can now be set by summing a correspondingly large slice of the movie stack for each crystal. The latter process can be repeated such that sets with different integration times are available, which can be compared later on, or used for different steps of the pipeline.

**Supplementary Figure 3:**
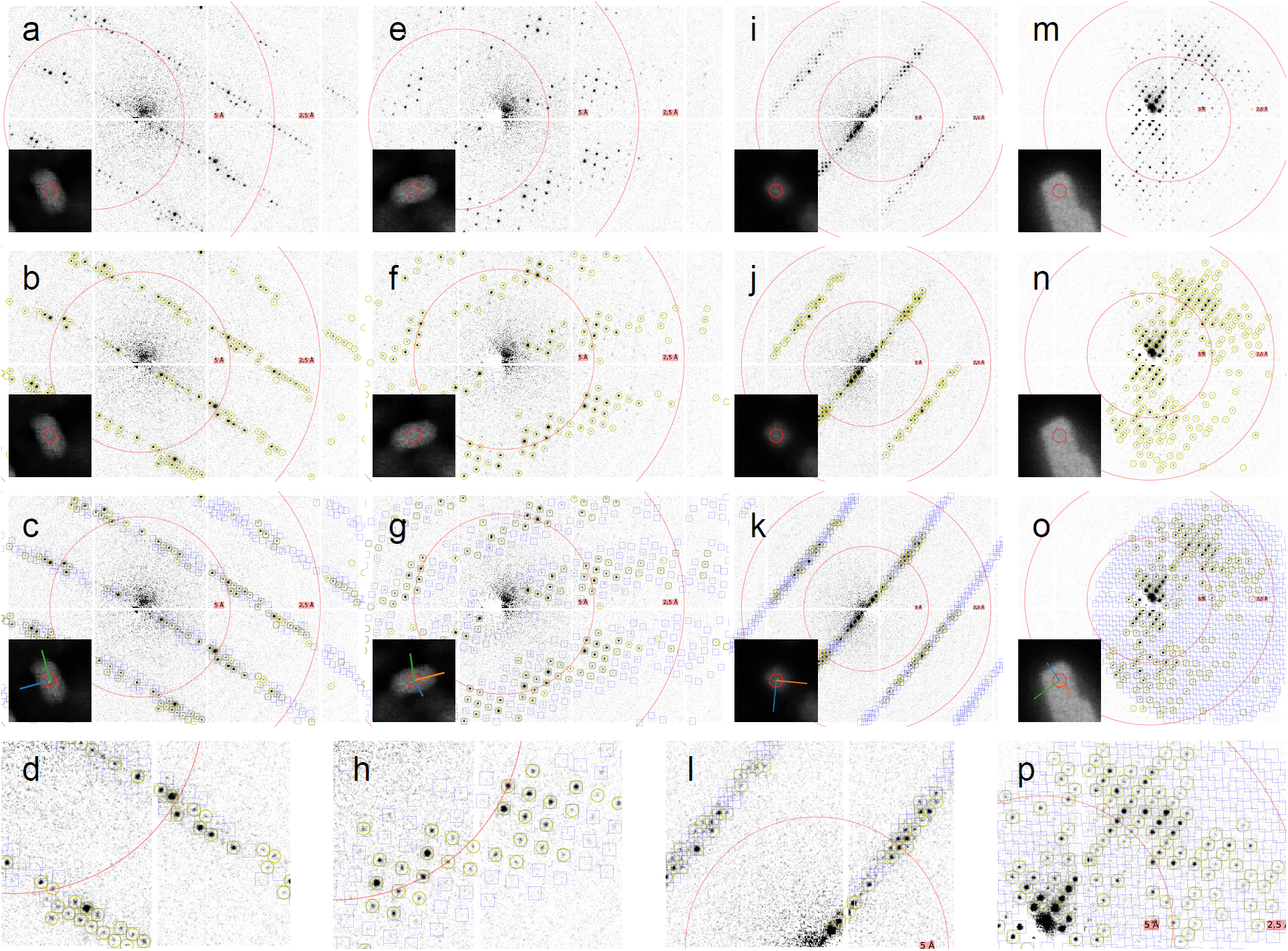
Data processing pipeline. (a) Diffraction pattern from a single granulovirus after dead-pixel and flatfield correction, and background subtraction. As the diffraction pattern is not centred of the detector, the resolution rings shown at 2.5 Å and 5 Å are not aligned with the pattern. The inset shows a close-up of the STEM mapping image on the corresponding granulovirus; the red circle corresponds to the beam diameter of 100 nm. (b) Same pattern after determination of beam centre and diffraction peaks, marked in yellow. (c) Same pattern including predictions of Bragg reflection positions after running indexing and refinement in *PinkIndexer*, shown as blue squares. The lines in the mapping image inset correspond to the derived real-space lattice vectors. For single granuloviruses it is typically found, that one of the lattice vectors is aligned with the long axis of the virus shell. (d) Zoom into a region of (c), highlighting the matching between predicted peaks (blue squares) and pixel intensity data. (e-h) As (a-d), for another virus. (i-p) As (a-h), for two lysozyme nanocrystals.
- Dead-pixel correction is applied by either replacing all dead pixels with a given integer number (typically −1 or NaN) or interpolating from adjacent pixels. Optionally, flat-field or detector saturation corrections can be applied by multiplying each pixel value with a previously determined normalization value, which can itself be a polynomial function of the pixel value. The pixels near the gaps of the 12 detector panels, which are three times more elongated in the direction facing the gap and hence have a different effective gain and saturation behaviour, can either be omitted from the analysis, or scaled to have their intensity matched with the other pixels. In the present work, we chose to omit these pixels.
- The centre of each diffraction pattern is determined in a multi-step process, and the images are correspondingly shifted (Supplementary Figure 3, second row). This is mandatory, as even for a good alignment of the STEM pivot point before data acquisition, a slight position-dependent beam tilt will remain. This manifests as displacement of the diffraction pattern, hampering the accuracy of the subsequent indexing step. First, the centre-of-mass of pixel intensities within the inner region of the image is found for each shot. Next, the obtained position is used as a starting value for least-squares fitting of a rotationally symmetric Lorentzian function over a small domain (30×30 pixels) around the centre-of-mass position. Finally, if peaks are found in the diffraction pattern, a refinement of the centre position is performed by matching the position of Friedel-mate reflections, which are generally found at low resolutions. Further refinement of the centre of each diffraction pattern is done at the indexing step (see below).
- Optionally, the radially symmetric background in the diffraction patterns, which is caused by inelastic scattering events that do not contribute to Bragg peaks, can be subtracted. This is done by azimuthal integration at each radial pixel coordinate, whereby regions around each Bragg peak are ignored. The derived radial profiles are then median-filtered and subtracted from the images.

The final result of this pipeline is a data stack containing the corrected, dose-integrated and centred diffraction data and corresponding metadata of all diffraction shots, contained in *NeXus*-compatible HDF5 files. We could successfully export the data to the *CrystFEL* ^57,59^, *DIALS* ^70^, and *nXDS* ^71^ packages.

### Data reduction

To obtain a fully merged crystallographic data set from the single-crystal snapshots, we use the tools provided in *CrystFEL* 0.8.0 ^57,59,72^. Bragg reflections in the diffraction patterns are registered using the *peakfinder8* algorithm ^58^ (Supplementary Figure 3, second row). Because this algorithm internally estimates the radially symmetric background for each resolution shell, we have found no increase of accuracy when background subtraction is applied to the diffraction patterns before peak finding. As the first frame of each dose-fractionation movie may still contain slight artefacts arising from residual beam motion (Figure 3b), the reliability of peak finding can be increased by applying it to images summed from the stacks such that the first frame is excluded. Before the peak integration step, the first frame can be included again.

### Indexing and integration

One of the most difficult tasks when processing a single electron diffraction pattern is to find the orientation of the crystal that generated this pattern. Due to the very short de Broglie wavelength of electrons (0.025 Å at 200 kV, as compared to several Å in the case of X-rays), the measured part of the Ewald sphere is almost flat in the resolution range used for the measurements. Therefore, hardly any three-dimensional information can be extracted from a single pattern. To overcome this limitation, prior unit-cell information has to be used as a constraint, as it is done in various indexing algorithms, such as *TakeTwo* ^73^, *FELIX* ^74^, *problematic* ^75^, *SPIND* ^76^, or *PinkIndexer* ^39^. Having tested various algorithms, we found *PinkIndexer*, which can be used as a part of the *CrystFEL* package, to achieve the highest indexing rates for our data, at reasonable performance (roughly 30 seconds per pattern and CPU core at sufficiently fine sampling settings). Patterns containing too many observed diffraction peaks which cannot be assigned to any Bragg reflection (as predicted by the indexing result) indicate the presence of multiple crystals in that shot and are rejected. After successful indexing, the integrated image intensity is determined near each predicted Bragg peak position by background-subtracted summation (*rings-nocen-nograd* integration option in *CrystFEL*) ^72^. We found that despite the basic background correction that is applied when integrating each Bragg reflection, a global background subtraction as described above leads to significantly improved outcomes in this integration step.

### Merging

The data set is then merged using *partialator* ^59^, yielding a plain-text *hkl*-File containing the full reduced data set. Post-refinement and partiality modelling algorithms currently available in *partialator* have been found to be ineffective for our data; this will be investigated further in future work.

## Supplementary Discussion

In Supplementary Figure 4, two-dimensional cross sections through reciprocal space with Bragg spots of hue corresponding to the number of observations in the data sets are shown. For the lysozyme data set, reflections away from the symmetry axes are disfavoured, clearly indicating a preferred orientation with the facets of the tetragonal crystals parallel to the support film. The completeness of the data set is thus reduced to 78% up to 1.8 Å resolution. We note that this could have been mitigated by varying the sample grid tilt angle between regions; unfortunately, this was not performed here due to technical issues with the microscope during the time of the measurement. For the granulovirus data set, where, on top of the higher point group symmetry, tilt angles were varied and more crystals were recorded, complete coverage is achieved.

**Supplementary Figure 4:**
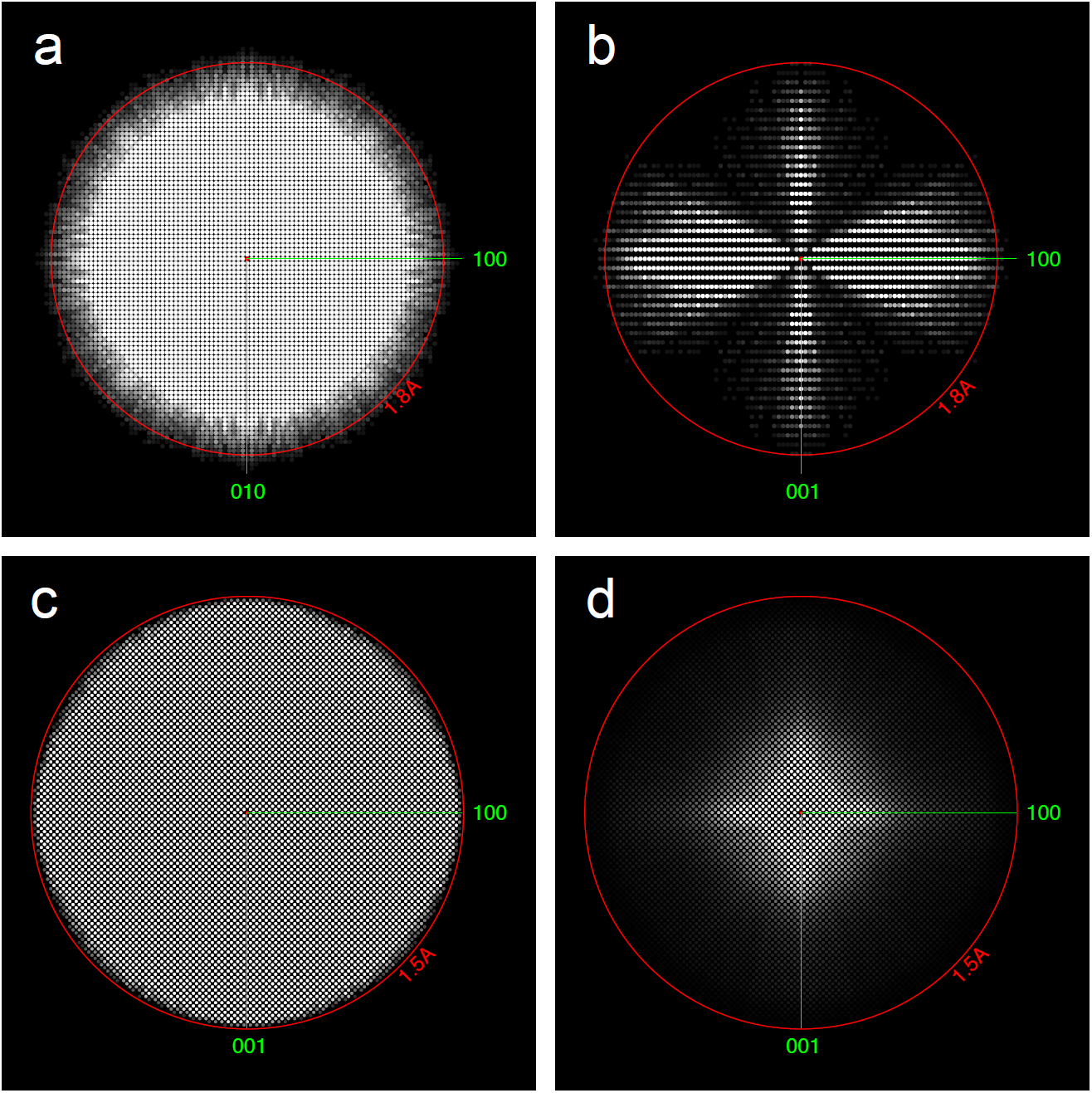
Data set completeness. (a, b) Number of unique Bragg-reflection observations in the lysozyme data set, shown in the (a) l = 0 and (b) k = 0 planes. White points correspond to on average 5 or more observations of each symmetry-related reflection. The red circle indicates a resolution of 1.8 Å. (c, d) Granulovirus data set, in l = 0 plane with different hue scale; white points correspond to (c) 5 or (d) 200 or more observations of each symmetry-related reflection. The red circles indicate a resolution of 1.5 Å.

